# Neural geometry efficiently representing abstract form of value and modality in the primate basal ganglia

**DOI:** 10.1101/2024.09.22.614265

**Authors:** Seong-Hwan Hwang, Ji-Woo Lee, Sung-Phil Kim, Hyoung F. Kim

## Abstract

Basal ganglia process diverse values from various modalities using limited resources, necessitating efficient processing. This involves converging tactile and visual information within bimodal value-coding neurons, ensuring efficient processing with limited number of neurons. However, convergence at the single neuron level may compromise modality-specific information, raising the question: Does efficient processing inevitably degrade information quality? We investigated the representational geometry in the putamen of macaque monkeys trained to learn values from tactile and visual inputs. Here, we demonstrated that the population representation of bimodal value-coding neurons in the putamen preserved both value and modality information, and these representations were shared to efficiently maintain quality. Notably, these representations were generalized across identical modalities and values, resulting in an efficient low-dimensional representation. Furthermore, a faster transformation to a generalized value representation within neural geometry reflected greater confidence in value-guided choice behavior—this correlation not observed in conventional decoding. Our results suggest that bimodal value-coding neurons play a key role in balancing efficiency and information fidelity, facilitating cognitive states required for confident decision-making.

## Introduction

Our brain efficiently processes numerous functions with a limited number of neurons, distinguishing it from contemporary artificial intelligence, which can expand its computing resources. This constraint is especially pronounced within cortico-basal ganglia circuits, which undergo a notable reduction in neuron count due to spatial limitations^1–3^. This anatomical arrangement necessitates striatal neurons to process multiple cortical inputs in a convergent manner^1–6^. Interestingly, we have demonstrated this convergent processing, particularly in integrating value information across different modalities at the single-neuron level in the primate putamen^7^. Bimodal value neurons, which encode both tactile and visual value information, constituted more than half of the value-coding neurons in the putamen, underscoring the striatum’s ability to efficiently handle a variety of functions with fewer neurons. This indicates that the striatum employs an efficient information processing strategy in a quantitative manner.

Pursuing quantitative efficiency alone does not guarantee information quality; instead it often compromises the quality. In our previous study, if bimodal value neurons integrate tactile and visual value information into a unified value representation without preserving their modality features at the single-neuron level, it potentially leads to a decline in information quality^7^. This issue is critical, as it may result in the failure to transmit modality information to subsequent neural layers in the basal ganglia structures, causing failures in decision-making due to inaccurate information flow. Therefore, preserving the distinctiveness of information while processing multiple streams is important for ensuring accurate information flow and effective decision-making. The brain should have mechanisms to ensure both quantitative efficiency and qualitative preservation of information. A plausible strategy for achieving both is by processing modality information at the neural population level in the putamen. Using value convergence at the single-neuron level and encoding modalities at the population-neuron level could enhance the efficient process of value and modality information even with a decreased number of neurons across the cortico-basal ganglia circuits.

Another aspect of efficient information processing involves the ability to generalize multiple pieces of information based on their shared common features. Structured neural representations, leveraging these shared features, may enable brain regions to process information more concisely and efficiently, using fewer variables. This generalization strategy allows for the processing of diverse information within lower dimensions in neural representations, as observed in the prefrontal cortex and hippocampus of primates^8^. Rather than requiring every neuron to examine every detail of individual pieces of information, processing based on shared features offers potential efficiency gains. Considering energy consumption, generating a greater number of neural patterns requires more energy because transitioning the brain states necessitate synaptic changes, such as alterations in ion channel responses and structural modifications in synapses, all of which demand ATP supply. Instead of requiring every neuron to examine every detail of individual pieces of information, processing based on shared features offers potential efficiency gains by saving energy needed for state changes. The capacity for generalization with geometrical neural representation appears to be linked with systematic process that facilitates its efficiency in managing diverse types of information. While low-dimensional representations have primarily been identified in cortical areas and the hippocampus, regions associated with high cognitive functions, it remains unclear whether the similar representations exist in primate basal ganglia structures interconnected with cortical areas^8–11^.

Moreover, the cognitive ability to generalize existing knowledge is crucial for efficiently guiding goal-directed behavior, as it enables rapid learning and focuses on shared features^12,13^. Neural processing also represents this generalization, with changes in neural geometry reflecting variations in behavior and cognitive states^8,9^. These findings suggest a link between goal-directed behavior and the capacity to generalize information, with corresponding geometric alteration in neural representations. Given the role of the basal ganglia system in processing value to guide goal-directed behavior, we aim to investigate how dynamic changes in neural geometry for value and modality within the putamen relate to behavioral outcomes^14–17^.

In this study, we examined neural geometry in the primate putamen, analyzing representations for each modality and value through cross-condition generalization performance (CCGP)^8^. Our findings revealed that bimodal value neurons efficiently encoded both modality and value with low-dimensional representation, exhibiting dynamic changes in their geometric structures across task periods. Furthermore, we conducted analyses to investigate whether these dynamic changes in neural geometry were correlated with cognitive states during the execution of value-guided behavior.

## Results

### Convergent processing of tactile and visual values in the primate putamen

To examine how population of neurons in the primate putamen processes value information from tactile and visual inputs, we trained monkeys to perform both tactile and visual modality-value reversal tasks as previously described (Fig. 1A)^7^. Data used in this study were from our previous study. In tactile and visual value reversal tasks (T-VRT and V-VRT), one braille pattern or fractal image was associated with reward (good), and the other was not (bad). This stimulus-reward contingency was reversed after 50 trials, enabling the examination of neural responses encoding the value while excluding neural responses to the stimuli (Fig. 1B).

**Figure 1.**
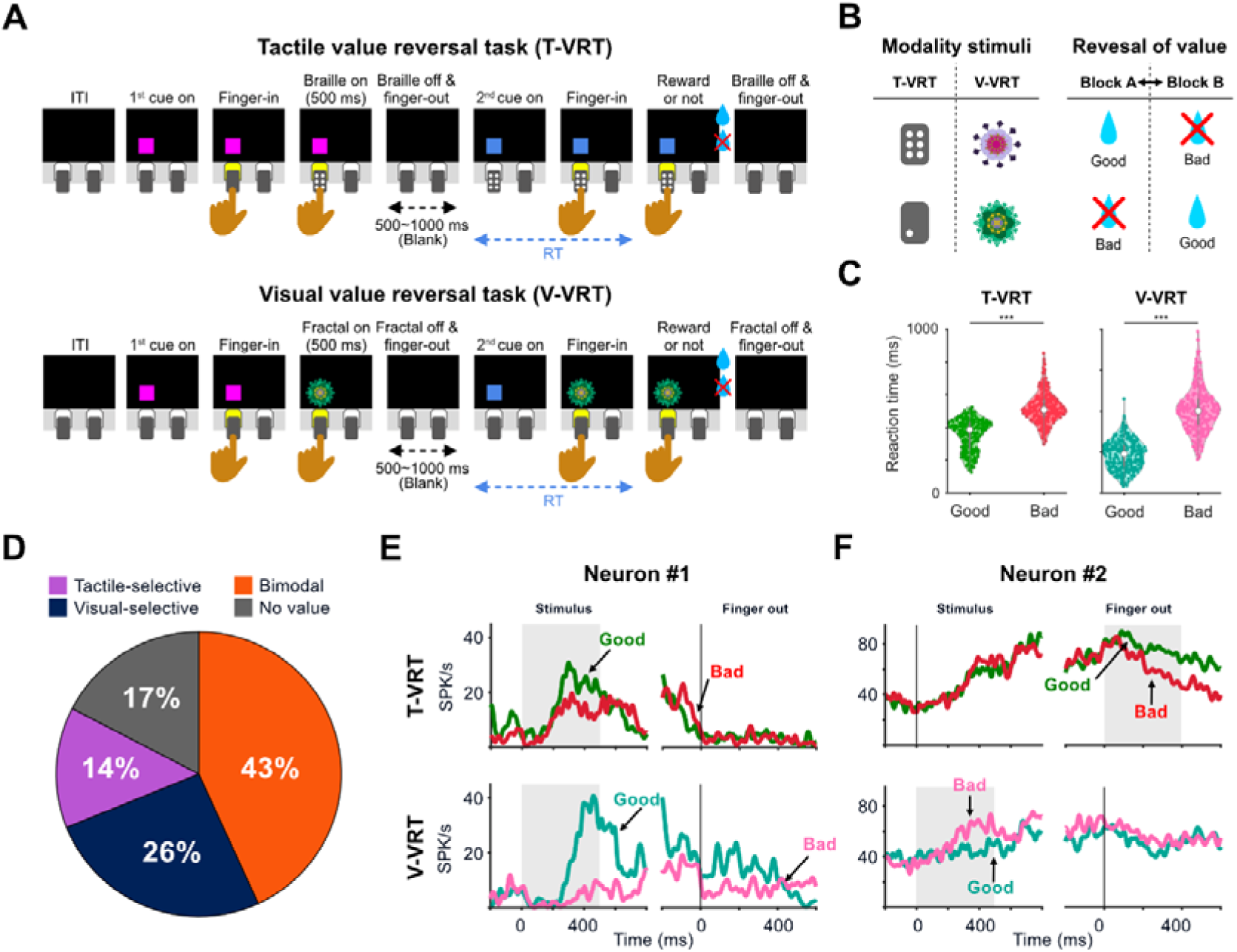
Tactile and visual value-guided finger insertion behavior and bimodal value neurons in the primate putamen. (A) Procedures of two different modality value reversal tasks: tactile value reversal task (TVRT) and visual value reversal task (V-VRT). The two tasks are identical in procedure, with the only difference being the type of stimuli presented. (B) In T-VRT, braille patterns were used as tactile stimuli. In V-VRT, fractal images were used as visual stimuli. To avoid the confusion in neural responses to the stimulus and to value, reversal of value was applied. (C) Differences in reaction times for finger insertion after the onset of 2nd cue based on the stimulus’s associated value (n = 299 for both the T-VRT and V-VRT). (D) Pie chart of recorded neurons in the putamen (n = 299). (E) Responses of an example bimodal value neurons in the primate putamen neuron in the putamen in T-VRT and V-VRT. (F) Responses of another example bimodal value neuron.

In both tasks, the same tactile or visual stimulus was presented twice after presentation of a square cue indicating finger insertion, enabling the monkeys to experience the stimulus during the 1^st^ cue presentation (stimulus presentation period) and predict the reward outcome during the blank delay period before the 2^nd^ cue presentation (Fig. 1A). Consequently, the monkeys inserted their finger faster into the hole after the 2^nd^ cue presentation when the previously experienced good stimulus was presented compared to bad stimulus (paired t-test, p < .0005 in T-VRT; p < .0005 in V-VRT) (Fig. 1C). The difference in reaction time indicates that the monkeys acquired and retained the values associated with the tactile and visual stimuli until the 2^nd^ cue presentation.

Three types of value-coding neurons were identified in the primate putamen through single-unit recording, as previously reported: tactile-selective, visual-selective, and bimodal value neurons (Fig. 1D)^7^. Bimodal value neurons comprise 43% of all task-related responsive neurons (129/299) as well as 52% of all value-coding neurons (129/247) (Fig. 1D).

Notably, we observed that bimodal value neurons in the putamen showed value discrimination responses in both T-VRT and V-VRT, but often during different periods (Figs. 1E and F). Figure 1E shows an example neuron that exhibited stronger responses to good tactile or visual stimuli compared to bad ones during the stimulus presentation in both T-VRT and V-VRT. However, this response for encoding both values was often not straightforward (Table S1). For instance, a neuron in Figure 1F encoded both tactile and visual values, but it represented each value during different periods: visual value was encoded during the stimulus presentation period while tactile value was encoded during the blank delay period. Taken together, although over half of value-coding neurons encodes both tactile and visual value information, the way to encode that information has heterogeneous response profiles across different task periods.

### The population of bimodal value neurons represents tactile and visual values differently

This diversity in the value encoding of bimodal value neurons raises a question (Fig. 2A): Do the population responses of these neurons integrate value information from tactile and visual inputs into a unified value, or do they separately represent the tactile and visual values in their population responses?

**Figure 2.**
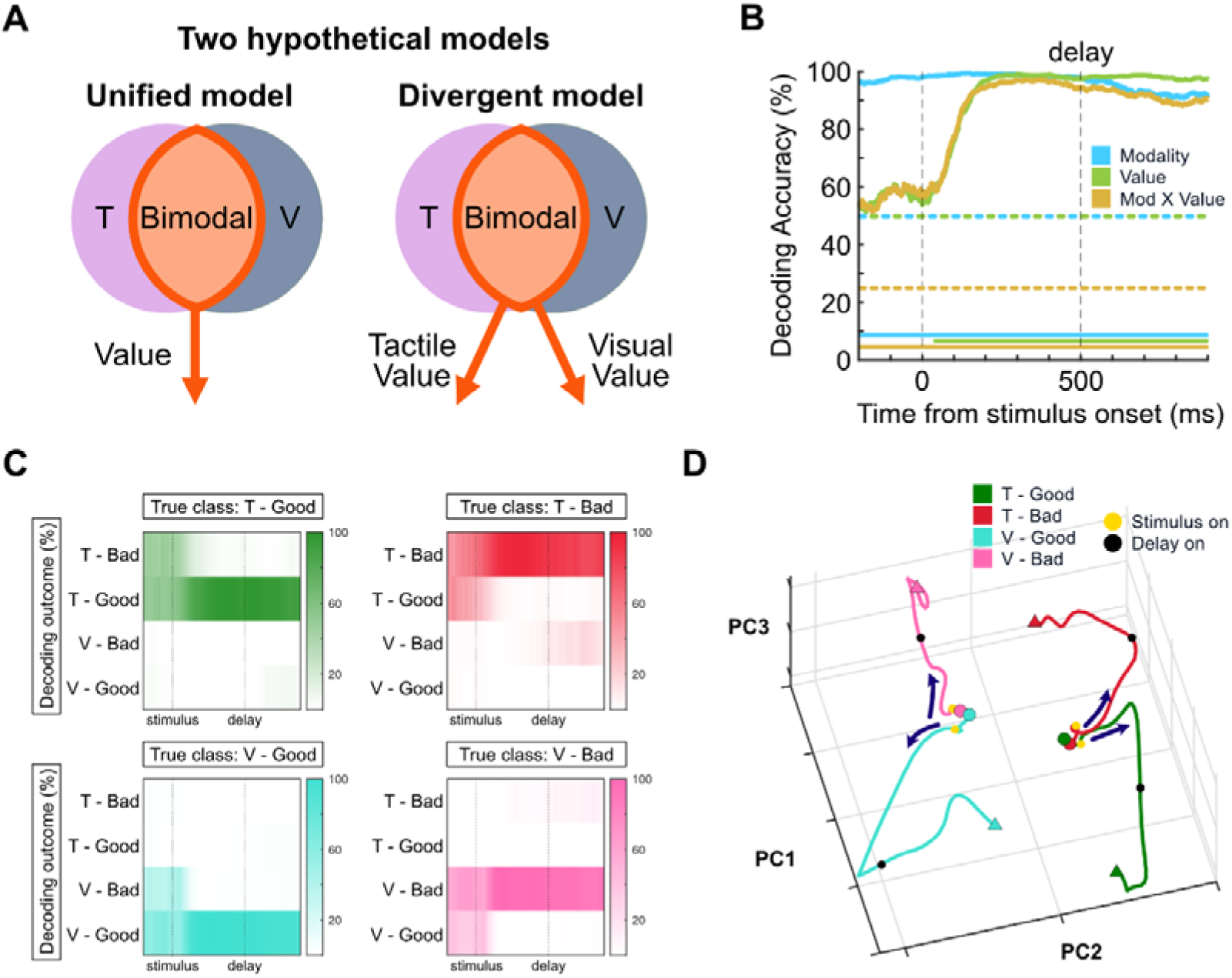
The neural ensemble of bimodal value neurons in the putamen represents both modality and value information. **(A)** Illustration of two hypotheses of the way bimodal value neurons process both modality and value information. **(B)** Decoding performance plotted as a function of time for each variable in bimodal value neurons: modality, value, and interaction of modality and value. Horizontal bars represent periods of decoding accuracy above the permutation results (right-tailed z-test, p < .05). **(C)** Confusion matrix for four different conditions. Each graph has a different class variable and shows responses of the decoder. **(D)** An example neural trajectories of bimodal value neurons (Monkey EV) in the subspace projected to the first three principal components (PCs). The circle and triangle denote the start (-200ms from the stimulus onset) and end points (1000ms from the stimulus onset) of neural trajectories, respectively. Yellow and dark circles indicate the stimulus on and delay on, respectively.

Previous studies have suggested that complex neural activities may serve latent functions involved in processing various types of information, which can be elucidated through population-level analyses^18–21^. Therefore, to determine whether bimodal value neurons process each value separately or in a unified manner, we conducted population decoding analyses of bimodal value neurons on three variables: modality, value, and their interaction (Figs. 2B and S1A).

The decoding accuracy for value was at chance level (50 %) before the onset of stimulus, but it quickly started to increase as the stimulus was presented. In contrast, modality was decoded nearly 100% prior to the stimulus onset, suggesting that the bimodal value neurons process modality information akin to contextual information in each task. The decoding accuracy for the interaction between value and modality (Tactile-good / Tactile-bad / Visual-good / Visual-bad) also increased and reached close to 100% after stimulus presentation, but the accuracy was already around 50% even before stimulus presentation (chance level = 25%). Considering that the bimodal value neurons represented the modality from the beginning of the trials, this statistically significant decoding for interaction prior to stimulus onset might be due to the use of modality information. To test this possibility, we constructed a confusion matrix of the decoders testing interaction between value and modality. The results indicated that the decoder differentiated between tactile and visual modalities prior to stimulus onset, and commenced discriminating values specific to each modality condition after stimulus presentation (Fig. 2C). These results indicated that the bimodal value neurons process modality as contextual information and discriminate values selectively in each modality condition.

These neural representations of modality and value dynamically changed over time (Fig. 2D). We compared the dynamics of the population activity among four conditions representing the combinations of modality and value in the subspace projected to the first three principal components (PCs) (Figs. 2D and S1B-C). Consistent with the decoding results, the trajectories revealed that the patterns of dynamics in latent space initially clustered by modality and then diverged according to their value and modality after the stimulus presentation. Overall, our data indicate that bimodal value neurons represented value differently depending on its modality, suggesting that the neural population in the putamen maintained the unique feature of modality when processing value components.

### Dynamics of the geometry of value and modality abstraction in the putamen

This distinct process of tactile and visual values at the population level of bimodal value neurons raises a question: Do the value and modality information form structured representation in bimodal value neurons? Can the representation of value and modality be generalized with shared feature? If the multiple variables can be generalized with a shared feature, the neural geometry of them forms an ‘abstract representation’, allowing for a reduction in neural dimensions when processing multiple inputs^8,9,12^.

To address these questions, we measured how well each variable is generalized using the cross-condition generalization performance (CCGP), as previously reported^8^. In the CCGP analysis, we determined whether a decoder, trained to identify the value (or modality) in one of two conditions (e.g. good and bad in tactile condition), could decode the same value (or modality) in a condition not used for training the decoder (e.g. good and bad in visual condition) (Fig. 3A).

**Figure 3.**
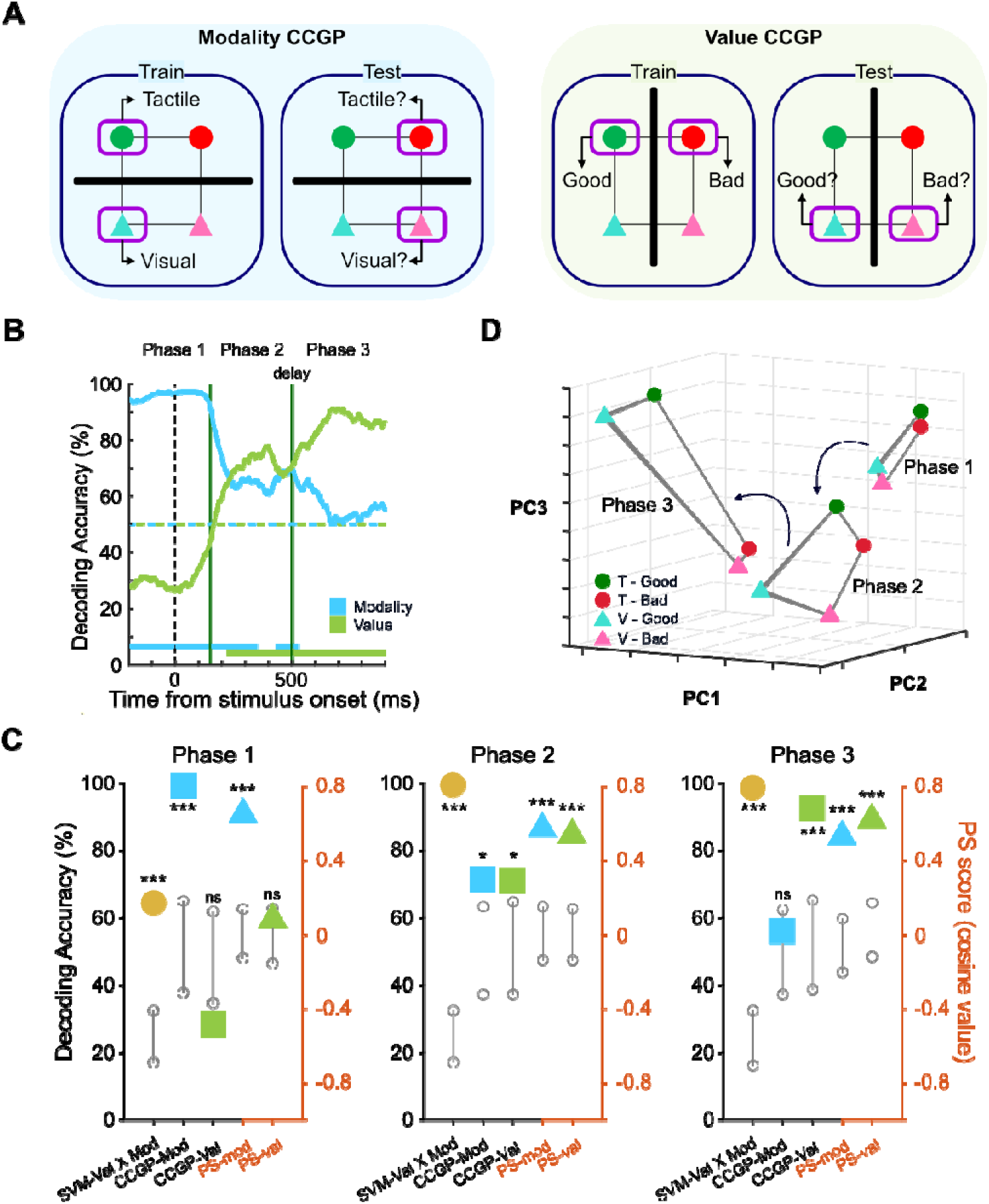
Dynamic change in the generalized representation of bimodal value neurons in the putamen over time. **(A)** CCGP scheme for modality and value. Green and red circles indicate tactile good and tactile bad conditions, respectively. Cyan and pink triangles indicate visual good and visual bad conditions, respectively. Different conditions were trained and tested to measure the extent of generalization of each variable. **(B)** CCGP plotted as a function of time for value and modality information in bimodal value neurons. Horizontal bars represent periods of decoding performance above the results of null model (right-tailed z-test, p < .05). **(C)** Comprehensive results of three analytic approaches (traditional decoding using SVM, CCGP, and PS) in three different time phases. The colored indicators represent the mean of each result, while the grayscale indicates the 95% confidence interval of null model. p values driven from Z score of data compared with null model (right-tailed for SVM and CCGP, twotailed for PS). *p < .05, **p < .005, ***p < .0005. **(D)** PCA plot depicting the representational geometry of bimodal value neurons over time with respect to value and modality.

Figure 3B illustrates the dynamic change of the CCGPs for value and modality as the trial progresses, aligned with the stimulus presentation (see also Fig. S2A-B). The CCGP for modality was initially above the chance level and sustained until about 150ms after the stimulus presentation. In contrast, the CCGP for value increased after the stimulus onset, surpassing chance level 230 ms after the stimulus appeared. The CCGP for value continued to rise after the stimulus onset and during delay period, while the CCGP for modality continued to decline during the stimulus onset and at the beginning of delay period.

Overall, the CCGPs for value and modality were initially distinct, then became similar, and eventually reversed as the trial progressed. To provide clear verification and visualization of this observation, we divided the time window of trial into three distinct phases: Phase 1, which encompasses pre-stimulus periods, from -200 to 150 ms aligned with the stimulus onset; Phase 2, focusing on stimulus presentation, from 150 to 500 ms; and Phase 3, covering the blank delay period, from 500 to 1000 ms (For further details and rationale behind this segmentation, please refer to the methods section). We further analyzed these three phases by determining which variables (value and modality) were generalized across shared features in each phase (Figs. 3C and S2C). In phase 1, the CCGP for modality was above chance level, but for value it was not (Fig. 3C, left panel). In both phase 2 and 3, the decoding accuracy for the interaction between value and modality was above 98%, as analyzed by a traditional linear decoder (Fig. 3C, middle and right panels). However, the extent of generalized representation dynamically changed as the phase progressed. In phase 2, CCGPs for both value and modality were above chance level (Fig. 3C, middle panel). However, in phase 3, the extent of generalized representation for value further increased, but the extent of generalized representation for modality decreased at the chance level, as analyzed by CCGP (Fig. 3C, right panel).

To more clarify the representational geometry of bimodal value neurons, we further computed the parallelism score (PS) in each phase, as previously reported, to quantify the degree to which coding directions are parallel^8^. If the coding vectors for each variable are nearly parallel, the parallelism score (PS) will deviate significantly from 0. Conversely, if the neural representations of each variable are similar to random representations, the PS will approximately be 0, indicating orthogonality between the coding vectors. As an example from our actual experiment, we can obtain coding vectors for classifiers trained to classify values as tactile-good and tactile-bad, as well as classifiers trained to classify values as visual-good and visual-bad. We can then calculate the cosine angle between these two coding vectors and define it as the PS for ‘value’. If the cosine angle between these two coding vectors is close to 1, it indicates that they are parallel.

As the result depicted in Figure 3C, the PSs for value and modality in all phases were above the chance level, except for the PS for values in phase 1. This suggests that the coding direction for both modality and value are almost parallel to the coding direction for that same variable across conditions after stimulus onset, potentially leading to a high decoding accuracy and CCGP in phase 2 and 3. Taken together, our quantification analyses using CCGP revealed that the generalized representation for modality decreased, while the generalized representation for value increased as decision-making progressed.

To visualize the geometric architecture of these neural representations, the average population activities for all four possible pairings of value and modality in each phase were projected into a three-dimensional PC space (Fig. 3D and Movie S1). In phase 1, the population activities clustered according to the modality, showing the similar representation across same modality. Interestingly, as they progressed to phase 2, each variable diverged, forming a square-shaped geometry that captures low dimensional representations for both value and modality. Subsequently, this geometric arrangement stretched along the value axis during delay period of phase 3, resulting in a rectangular-shaped geometry that captures the similar representation across same value, but not modality. Our comprehensive analytical approaches, including CCGP and population responses in latent dimensions, successfully demonstrated that bimodal value neurons form low-dimensional representations. Furthermore, this neural geometry exhibits dynamic changes as trial progressed, primarily stretching out along the value axis.

### Representational geometry for value and modality is related with the performance of value-guided behavior

The ensemble of bimodal value neurons in the putamen dynamically shifted towards enhancing the generalized representation of value as value-guided behavior is imminent. We thus investigated whether this dynamic shift correlated with the performance of the value-guided behavior.

In our study, we assessed the performance of value-guided behavior by measuring reaction times of finger insertion. If monkeys could accurately recognize the value of the stimulus and anticipate the availability of a reward, they would exhibit either a quick or slow finger insertion behavior, indicating highly confident decision-making. In contrast, if monkeys were unsure about the stimulus value, they would not exhibit a clear difference in reaction times according to the values, indicating less confident decision-making. Thus, we divided the trials into two groups based on the reaction times of finger insertion: the high-confidence and low-confidence finger insertion groups (Fig. 4A). The high-confidence group included trials with faster reaction times to the good stimulus and slower reaction times to the bad stimulus, both in the upper and lower portions of the 50^th^ percentile, respectively. Conversely, the low-confidence group comprised trials with slower reaction times to the good stimulus and faster reaction times to the bad stimulus, in the lower and upper portions of the 50^th^ percentile, respectively.

**Figure 4.**
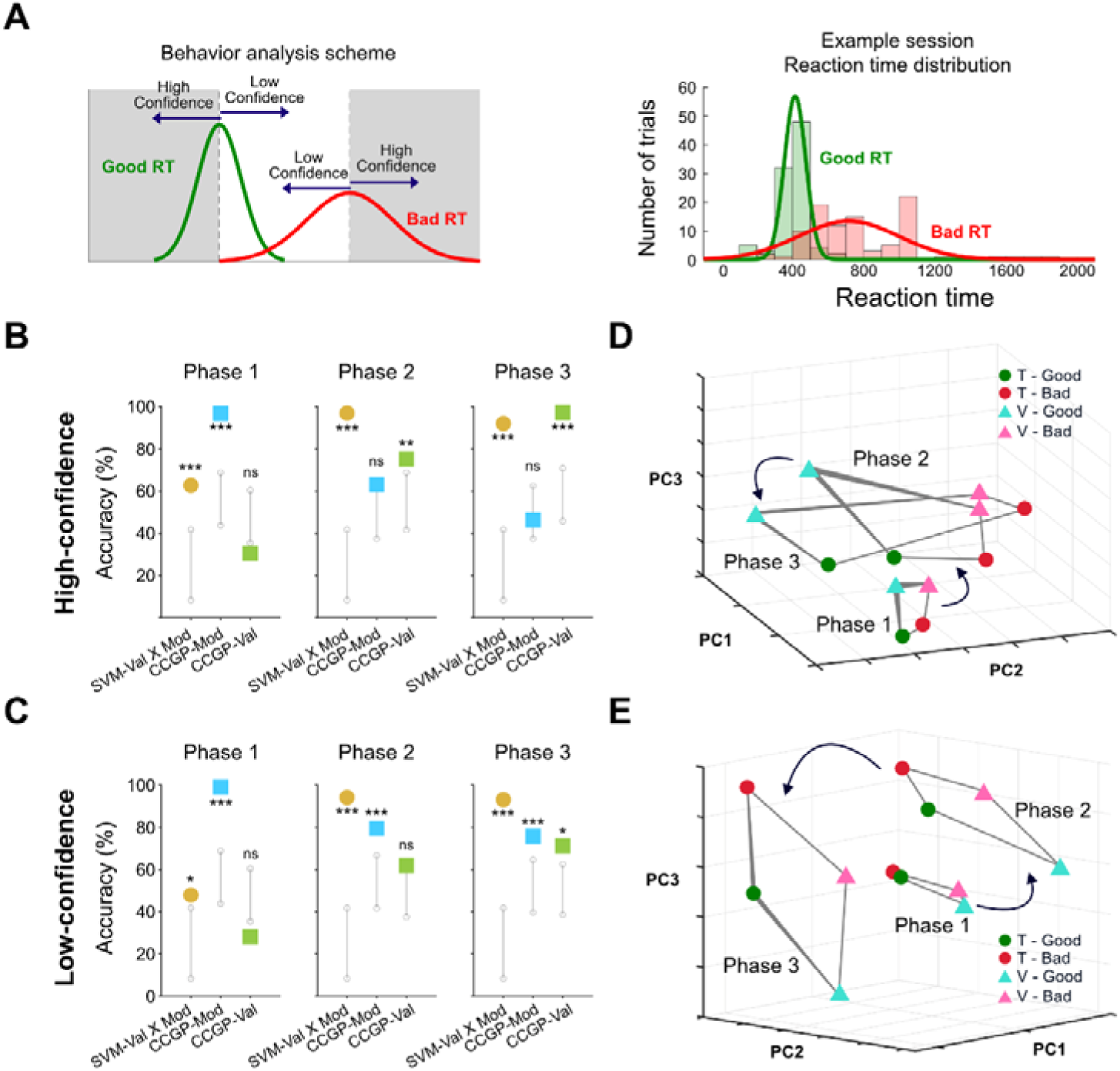
Correlation between behavior performance and the geometry of neural representation. **(A)** Analytic scheme for dividing trials to high- and low-confidence groups based on reaction times. The histogram of reaction times in actual example session is shown in the right panel. **(B)** Comprehensive results of traditional decoding (SVM) and CCGP in three different time phases with high-confidence trials. The colored indicators represent the mean of each result, while the grayscale indicates the 95% confidence interval of null model. **(C)** Similar to (B) but with low-confidence trials. **(D)** PCA plot illustrating the geometry of bimodal value neurons over time with high-confidence trials. **(E)** Similar to (D) but with low-confidence trials.

Traditional population decoding analysis for the interaction between modality and value showed no clear difference between the two groups across all phases. Particularly in phase 2 and 3, after monkeys experienced the value-associated stimulus, decoding accuracy approached nearly 100% in both groups (Fig. 4B and C). However, these two groups showed clear differences in generalized representations of value and modality as the decision of finger-insertion progressed (Fig. 4B and C). The CCGPs for modality were only statistically significant in both groups in phase 1. Subsequently, in phase 2 and 3, each group exhibited distinct changes. In the high-confidence group, the CCGP for modality dropped to the chance level in phase 2, while the CCGP for value increased above the chance level in phase 2 and reached nearly 100% accuracy in phase 3 (Fig. 4B). In contrast, the low-confidence group maintained the CCGP for modality above the chance level across all phases. Furthermore, the CCGP for value remained at the chance level by phase 2 and only slightly surpassed the chance level in phase 3 (Fig. 4C). Additional time-resolved decoding analyses confirmed that the difference between two groups are driven from the extent of generalization of modality and value, not from the traditional decoding performance (Fig. S3).

This difference between the high- and low-confidence groups can be visualized by projecting the average population activity for all possible pairings of value and modality into a three-dimensional (3D) latent space (Figs. 4D and 4E). In the low-confidence group, the population activity retained similar representation along the modality axis until phase 3 (Fig. 4E). However, the population activity of the high-confidence group expanded more rapidly along the value axis as the phases progressed compared to the activity of low-confidence group (Fig. 4D). Taken together, our results indicate that the performance of value-guided behavior is more closely linked to the shaping of neural geometry than the simple neural representation for value and modality.

### Distinct contribution of two types of bimodal value neurons in the generalized representation of value and modality

In our previous study, we identified two types of individual bimodal value neurons in the putamen based on the periods during which they displayed stronger responses to value: stimulus value neurons and delay value neurons^7^. To investigate the distinct role of these different types of single neurons in processing the value and modality at the population level, decoding analyses were exclusively performed using neurons from each group (Figs. 5A-B and S4).

**Figure 5.**
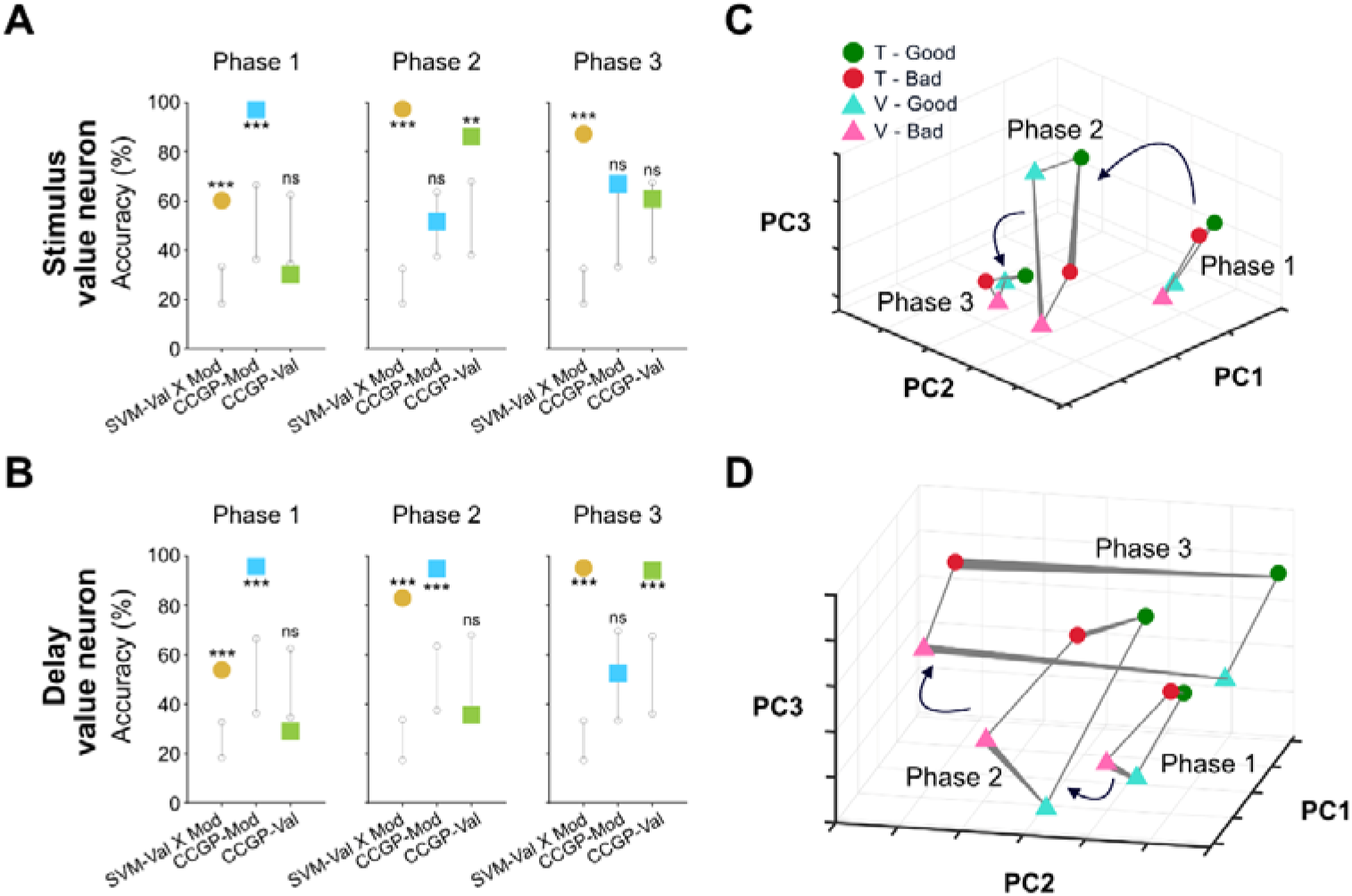
Subpopulation of neurons contribute different aspects of neural geometry. **(A)** Traditional decoding performance (SVM) and CCGP for stimulus value neurons. **(B)** Similar to (A) but with delay value neurons. **(C)** PCA plot depicting the representational geometry of stimulus value neurons with respect to value and modality. **(D)** Similar to (C) but with delay value neurons.

In traditional neural decoding, the decoding accuracy for the interaction between value and modality did not differ between the two neural types across all phases (Figs. 5A and B). This suggests that the contributions of both subpopulations were similar for processing the value and modality information. However, we found distinct dynamic changes of the generalized representation for value and modality between these two types of neurons through CCGP analysis.

In phase 1, both types of value-coding neurons exhibited high CCGPs for modality above the chance level, while the CCGPs for value were not statistically significant (Fig. 5A and B, left panels). In phase 2, the CCGP for value of the stimulus value neurons surpassed the chance level, while the CCGP for modality declined to the chance level (Fig. 5A, middle panel). Subsequently, in phase 3, their CCGPs for both value and modality were no longer statistically significant (Fig. 5A, right panel). Conversely, the CCGP for modality of the delay value neurons remained statistically significant in phase 2, while the CCGP for value was not (Fig. 5B, middle panel). In phase 3, their CCGPs for modality and value were reversed (Fig. 5B, right panel).

We also observed the distinct changes in geometries of each neural type projected into a 3D latent space (Fig. 5C and D). In phase 1, the population responses of stimulus value neurons were clustered according to the contextual modality, showing the generalized representation of modality (Fig. 5C). Their population responses were stretched along the value axis in phase 2, generating a rectangular-shaped geometry that captures the generalized representation of value. Finally, in phase 3, this geometry collapsed, resembling a random representation. In contrast, the population responses of delay value neurons retained their clustering according to the contextual modality until phase 2 (Fig. 5D). However, in phase 3, they diverged along the value axis, forming a rectangular-shaped geometry. Overall, both types of value neurons showed similar capabilities in conveying value and modality information, but they exhibited distinct dynamic changes in the neural representation of modality and value. This ablation study, excluding one of each neural type, elucidates that both neural types in the putamen were necessary for generating the factorized representation observed in phase 2, as depicted in Figure 3C.

### Selective generalized representation for modality but not value by modality-selective value neurons

Modality-selective value neurons, which encode either tactile or visual value selectively, constitute the remaining half of value-coding neurons identified in the putamen. We then examined the contributions of the modality-selective value neurons to the generalized representation of value and modality, as well as their correlation with value-guided behavior at the population level (Fig. 6A).

**Figure 6.**
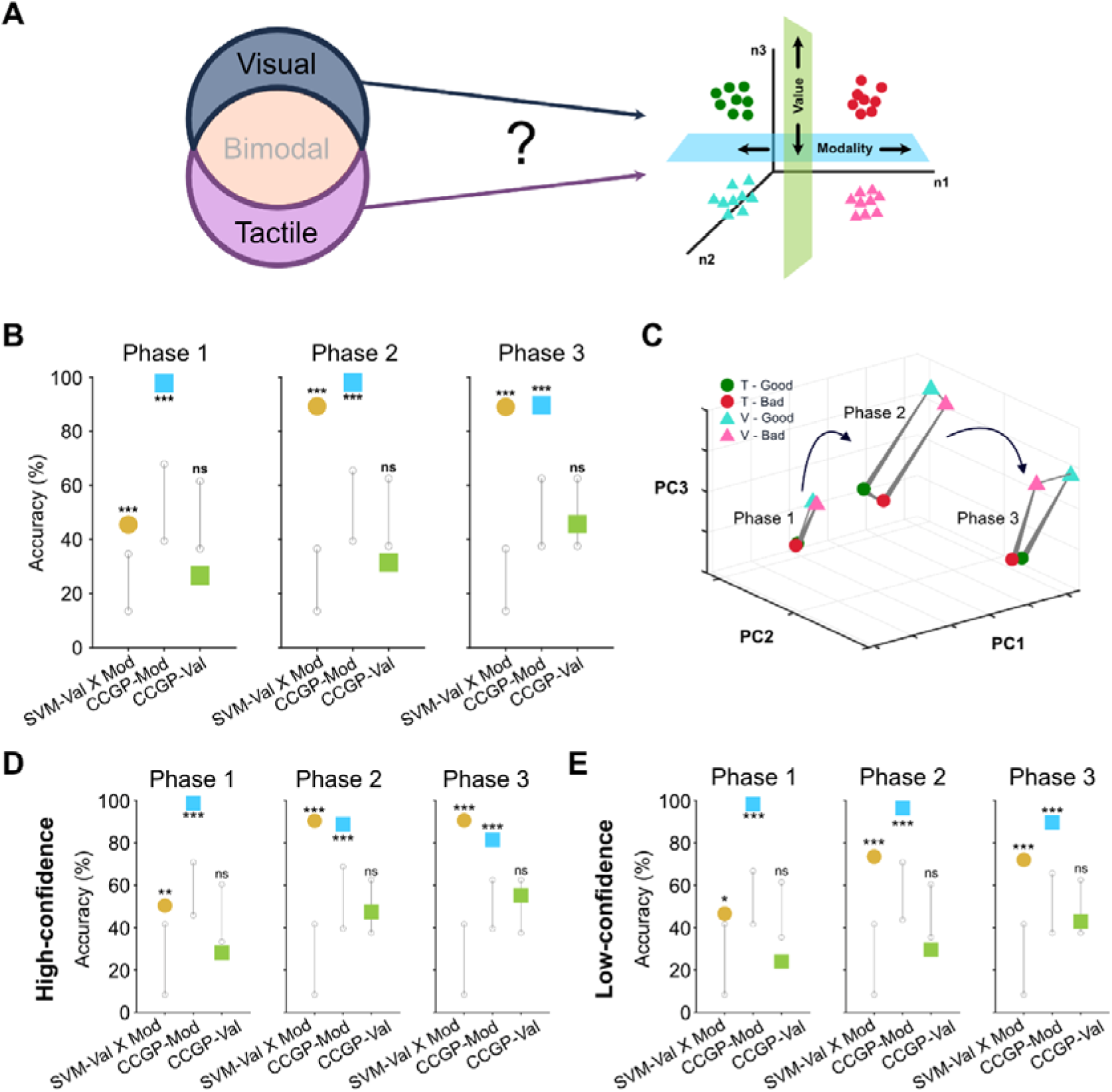
The selective role of modality-selective value neurons in generalization modality but not value. **(A)** Illustration of modality-selective value neurons when processing two different value information. We conducted the same analyses previously performed for bimodal value neurons using modality-selective value neurons. **(B)** Traditional decoding performance (SVM) and CCGP for modality-selective value neurons. **(C)** PCA plot depicting the geometric structure of modality-selective value neurons with respect to value and modality.

We found that the population activity of the modality-selective value neurons selectively represented the generalized form of modality, but not the value across all phases (Figs. 6B and S5A). In traditional neural decoding analysis, the modality-selective value neurons successfully discriminated between four variables in the phases 2 and 3, as demonstrated by the decoding accuracy for interaction (Fig. 6B, middle and right panels). Interestingly, their CCGPs for contextual modality were higher than the chance level, but not for value across all phases (Fig. 6B). We also visualized the dynamic change of neural representation by projecting them into the 3D latent space: the clustering according to the contextual modality was preserved across entire phases (Fig. 6C). Furthermore, there were no difference in dynamic changes of generalized representation between high and low-confidence groups (Figs. 6D-E and S5B-C). These findings suggest that the modality-selective value neurons play a selective role in generalizing modality information, rather than in generalizing value for value-guided behavior.

## Discussion

Our population analyses demonstrated that bimodal value neurons in the putamen, which converge value information regardless of modality inputs at single neuron level, preserve both modality and value information. Interestingly, the neural representation of these bimodal value neurons exhibited a highly organized geometric structure, showing similar representations across identical modalities and values. Furthermore, as a trial progressed, the representational geometry of these neurons underwent dynamic transformations. The dynamics of these changes reflected the level of confidence in finger insertion behavior. This relationship between changes in neural representation geometry and behavioral outcomes was not observed in modality-selective value neurons, indicating the unique role of bimodal value neurons in representing cognitive states that align with value-guided behavior. Our findings suggest that convergent processing in the primate basal ganglia system forms a low-dimensional population representation of values from various modality inputs, thereby efficiently preserving both value and modality information.

### Low-dimensional representation for efficient processing in the basal ganglia

How the basal ganglia structures process diverse information with limited neural resources has been a longstanding question in neuroscience^2,4–6^. Previous studies have suggested the presence of convergent processing in the basal ganglia due to its anatomical funneling structure, where the number of neurons decreased from the cortex to the output structures^3,5,7^. Moreover, it remains unknown how this convergent processing manages diverse information and transmits it to subsequent brain regions. Our results showed that bimodal value neurons in the putamen discriminated both value and modality information within their low-dimensional population representation (Fig. 3). This suggests that the basal ganglia system employs a strategy that achieves both objectives: quantitative efficiency by using fewer neurons while still efficiently retaining all the information at the population level.

It is noteworthy that putamen neurons process value and modality information using low-dimensional representations, a characteristic mostly reported in higher-order structures, such as the prefrontal cortex and hippocampus^8,11,22,23^. Is it then necessary for putamen neurons to process information as low-dimensional and generalized neural representations across the same modality or value features? If the sole purpose of the putamen is simply to relay information to subsequent structures, distinguishing and accurately conveying the information through population representation would be sufficient, without the need for a low-dimensional format.

One important point to consider is that the number of neurons decreases from the putamen to subsequent structures^24,25^. In this anatomical funneling structure, value information that is not efficiently processed could become a significant burden for the subsequent brain structures with fewer neurons, leading to improper value processing. Therefore, to convey information to subsequent structures without loss and to ensure proper processing in these downstream regions, the putamen might need to represent and transmit value information in a low-dimensional generalized form.

Additionally, although recent studies have discussed the advantage of low-dimensional representation for readout in downstream structures^23,26,27^, direct empirical evidence has not yet been demonstrated. It will be crucial to investigate the circuit-level mechanisms underlying low-dimensional representation of diverse values within basal ganglia structures in future research.

### Distinct roles of subpopulations of value neurons in the putamen

Understanding how single-cell properties impact population representation and how neural population patterns contribute to actual brain function remains a complex challenge in neuroscience^19,22,26,28–32^. Additionally, whether different populations of neurons perform distinct functions remains an ongoing question^33–36^. In our study, we identified different populations of neurons in the putamen based on their characteristic responses to value: stimulus and delay value neurons, as well as bimodal and modality-selective value neurons. This categorization allowed us to analyze the distinct roles of these subpopulations in processing value and modality.

We confirmed that all groups of value neurons were capable of distinguishing both value and modality information (Figs. 2b, 5, and 6). However, they exhibited significant differences in how and when they generalized modality and value information. By dividing bimodal value neurons into stimulus value neurons and delay value neurons based on their predominant value responses in distinct phases, we observed clear differences in the patterns and timing of generalized value information (Fig. 5). The differences were even more pronounced between bimodal and modality-selective value neurons (Fig. 6). While bimodal value neurons encoded generalized information for both modality and value, modality-selective value neurons exhibited generalized representation only for modality. Furthermore, bimodal value neurons demonstrated a relationship between the speed of emergence of the generalized neural representation for value and the confidence in value-guided finger insertion, whereas modality-selective value neurons did not (Fig. 6d-e). This suggests that each neural characteristic may contribute to different roles in shaping the geometry of representation rather than simply representing information.

### Dynamic transformation of neural geometry for generalized value representation and value-guided behavior

We found a close relationship between the extent of value generalization in the putamen and value-guided finger-in behavior, consistent with previous reports that demonstrate the link between abstract neural representation and behavior^8,9,37,38^. Notably, when monkeys exhibited highly confident value-guided behaviors, the generalized neural representation for value emerged much faster, and the neural geometric arrangement became more stretched along the value axis (Fig. 4). These findings raise a further question about what drives this stretching and the emergence of value-generalized neural representation.

One possibility is that this dynamic transformation of neural geometry is caused by a cognitive ability that allows subjects to focus on specific information. Fascianelli and colleagues showed that the variable exhibiting generalized representation in the prefrontal cortex varied depending on which aspect of the task each monkey focused on^37^. In our study, the monkey’s primary focus was whether or not a reward would be given in each trial. This suggests that the monkey’s cognition strategically prioritizes value information over modality information, which is naturally reflected in its neural representation. Therefore, when the monkeys focused more on value information, the neural representation in the putamen likely generalized more rapidly according to value, leading to clearer value-based behavioral differences.

This cognitive prioritization of value may be linked to strong inputs of the striatum from midbrain dopamine regions, which are involved in motivational control and value formation^14,15,34,39–43^. The dynamic changes in neural geometry observed in the putamen might be driven by dopamine neurons processing which information should be focused and which is most valuable. Understanding the role of dopamine neurons in shaping the neural geometry of the basal ganglia system will be crucial for uncovering the mechanisms underlying the formation and evolution of these geometries, as well as for predicting value-guided behavior.

## Materials and Methods

### General experimental procedures and subjects for electrophysiology

All of the data were collected in our previous study, and reanalyzed in this study ^7^. All experimental and animal care procedures were approved by the Seoul National University Institutional Animal Care and Use Committee.

Two rhesus macaques (*Macaca mulatta;* female monkey EV (5.2 kg) and male monkey UL (10.7 kg)) were used for the experiments. During the period of general anesthesia and surgical preparation, the monkeys’ skulls underwent implantation of a plastic head holder and a recording chamber. Each chamber was positioned at a lateral tilt of 25° to align with the putamen. Training and recording sessions commenced once the monkeys had completely recuperated from the surgery.

### Single-unit recording

During the task performance by the monkey, the activity of individual neurons within the putamen was recorded using standard procedures. Determination of recording sites utilized a grid system with 1mm spacing, aided by MR images (3T, Siemens) aligned with the chamber’s direction. Single-unit recording employed a glass-coated electrode (Alpha-Omega), inserted into the brain through a stainless-steel guide tube and advanced using an oil-driven micromanipulator (MO-974A, Narishige). Neuronal signals were amplified, filtered (250 Hz to 10 kHz), and digitized (30-kHz sampling rate, 16-bit A/D resolution) using a Scout system (Ripple Neuro, UT). Online isolation of neuronal spikes was achieved through custom voltage-time window discrimination software (BLIP, Laboratory of Sensorimotor Research, National Eye Institute National Institutes of Health [LSR/NEI/NIH], accessible at www.cocila.net/blip), with corresponding timings detected at 1 kHz. Individual spike waveforms were recorded at 50 kHz.

### Behavioral tasks

Monkeys performed two different modality value tasks, Tactile Value Reversal Task (T-VRT) and Visual Value Reversal Task (V-VRT) as previously described^7^.

In the T-VRT task, monkeys were trained to insert their left index finger into the left hole of a braille presentation case upon cueing by colored square cues on the screen. Once they inserted their fingers into the hole, a braille pattern was delivered and remained present for 500 milliseconds (ms). The monkeys were then instructed to maintain contact with the tactile stimulus until the disappearance of the finger-in cue and the tactile stimulus. Since the braille pattern was presented within an invisible braille case, monkeys relied solely on tactile perception to discriminate its value. After retracting their finger from the hole, there was a delay period ranging randomly from 500 to 1000ms. Subsequently, upon the onset of second cue, monkeys could either reinsert to touch the same stimulus that they had experienced before and receive a stimulus-associated reward, or give-up reinserting their finger by waiting 1000 to 2000ms. If they chose to reinsert the finger, they were required to maintain finger insertion for a minimum of 200ms to receive stimulus-associated reward. If they chose to give-up the reinsertion, then they would have to just wait for the next trial.

The reaction time for finger insertion was measured from the onset of the second ’finger-in’ cue to the actual finger insertion. Across two blocks, each consisting of 50-60 trials, the association between the stimulus and its value was reversed. Initially, in the first block, one stimulus was linked to a liquid reward, while the other was not. Subsequently, in the second block, this contingency was reversed. The order of the blocks was randomized across session, ensuring that the pairing of stimuli with rewards remained unknown to both monkeys and experimenters until the beginning of each session.

The procedure for V-VRT was identical as T-VRT, except for the presentation of visual stimuli instead of tactile stimuli. Fractal images, associated with actual reward, were displayed on the monitor screen when monkeys inserted their left index finger upon the presentation of a cue.

### Single neuron categorization

We initially categorized neurons exhibiting task-related responses. Evaluating the task-related neuronal responses involved isolating the activity of individual neurons and calculating the spike counts during T-VRT and V-VRT. For this assessment, we compared spike rates between the targeted test windows and control windows. Control windows were defined as the period 200–0ms before the 1^st^ finger-in cue on, while test windows spanned 0–500ms after each task event had begun. Task events included (1) 1^st^ finger-in cue-on, (2) 1^st^ finger-in, (3) Stimulus-on, (4) Stimulus-off, (5) 1^st^ finger-out, (6) 2^nd^ finger-in cue-on, (7) 2^nd^ finger-in, (8) Reward, (9) 2^nd^ finger-in cue-off, and (10) 2^nd^ finger-out. Statistical significance in task-related responses was determined by comparing spike counts between control and test windows in individual trials using the Wilcoxon rank-sum test. Neurons showing statistical significance in any of the tested task events were classified as task-related response neurons.

Among the neurons with task-related responses, we examined whether each neuron exhibited value discrimination responses during either the stimulus presentation in the 1^st^ cue period or delay period as previously described ^7^. We quantified the activity of each individual neuron in response to tactile or visual stimuli by calculating the spike counts within a specified test window (0-500ms after stimulus onset or 1^st^ finger-out) across individual trials. The firing rates of individual putamen neurons in response to good and bad value were compared using the Wilcoxon rank-sum test to determine the statistical significance of value discrimination. Subsequently, we analyzed the neural responses in both T-VRT and V-VRT. Neurons displaying distinct value discrimination responses exclusively during either of the two periods of T-VRT but not in V-VRT were categorized as ‘tactile-selective value-coding neuron’. Conversely, those demonstrating value discrimination responses solely during V-VRT but not in T-VRT were identified as ‘visual-selective value-coding neuron’. Neurons exhibiting value discrimination responses in both T-VRT and V-VRT were classified as ‘bimodal value-coding neuron’. The details for defining each type of value-coding neuron and the corresponding numbers are summarized in Table 1.

### Population decoding

We conducted pre-processing steps for population decoding. First, given our finding that the putamen encodes value responses in two distinct time phases-the stimulus presentation and blank delay period, we computed the spike train in both the stimulus presentation and the delay period. For the stimulus presentation period, we used a [-200, 500] ms window aligned with the stimulus onset, while for the delay period, a window [0, 500] ms aligned with the finger-out was used.

Subsequently, these spike trains from the two periods were concatenated. The reason we did not simply set the time window from [-200, 1000] ms from stimulus onset was because it would encompass finger-out movement between the stimulus being turned off and the start of the delay. As this study primarily focuses on value processing, we aimed to minimize neural responses caused by movements as much as possible by combining these two distinct time periods. Second, different analytic windows were used for either time-resolved decoding or fixed-window decoding methods. The time-resolved decoding for Figures 2B and 3B was done on spike counts measured in a 100ms moving window, with a 1ms step size. For fixed-window decoding, we summed spike counts in a fixed window corresponding to each time phase in Figures 3-6. We reported averaged decoding accuracy across iteration results and used it as the main dataset for analysis and visualization. Concatenated spike trains were then normalized using z-score to capture response changes across all neurons, rather than focusing solely on those with high firing rates.

Using these pre-processed activities of each neuron, we created pseudo-simultaneous population data. Trials from each neuron were randomly selected, and we then performed 10-fold cross-validation using a linear support vector machine (SVM) decoder to evaluate performance. This procedure was repeated over 100 resample runs, where different random pseudo-populations were created on each run. We presented decoding results for three task variables: 1) modality, 2) value, and 3) interaction between modality and value. We ensured that an equal number of trials per condition (for interaction between modality and value, tactile-good, tactile-bad, visual-good, visual-bad) was contributed by each cell included in the population decoding. For all analyses, data were aggregated across monkeys, since all key features of the dataset remained consistent between the two monkeys.

### Permutation model for testing significance of decoding performance

A permutation test was performed by shuffling the labels of each trial condition in each neuron. The same decoding procedure was used to generate a null distribution of the decoding accuracies. This procedure was repeated 1000 times and generated a full shuffled decoding null distribution for each time bin. We reported the theoretical chance level for each tested condition in the main figures and the 5^th^ to 95^th^ percentiles of the null distribution in the supplementary figures. The p-value was calculated through right-tailed z-test by comparing the decoder’s performance with the null distribution. If p-value were below .05, we defined the decoder’s performance was significantly higher than the permutation results. In addition, in order to maintain a conservative approach in our analysis when using time-resolved windows, we tabulated the consecutive significant points and reported significance only when at least 20 consecutive time points displayed significance.

### Dimensionality reduction

Dimensionality reduction was employed to visually represent the decoding outcomes and the structure of population neural patterns. This was achieved by conducting principal component analysis (PCA) on the averaged z-scored neural activities per condition. PCA was applied to a firing rate matrix with dimensions of CT X N, where C represents the number of conditions, T denotes the number of time points per condition, and N indicates the number of neurons. With the exception of Figure 2D, which depicted neural trajectories with time-resolved activities, we analyzed and visualized all neural data across both monkeys.

### The cross-condition generalization performance (CCGP)

To quantify the extent of generalization across shared conditions and the geometry of neural activities in population level, we conducted CCGP in our experiment ^8^. Unlike the traditional cross-validated decoding (SVM), where trials from all conditions are present in both train and test sets, CCGP used trials from distinct conditions for train and test sets. For instance, when testing for ’value’, we could designate conditions ’tactile good’ and ’tactile bad’ as the ’good’ and ’bad’ conditions for training a decoder (Fig. 3A). Subsequently, we assessed whether the trained decoder could accurately classify conditions ’visual good’ and ’visual bad’ as ’good’ and ’bad’, respectively. By evaluating the classification performance on data from entirely different conditions, we could assess the similarity of population neural patterns among distinct conditions with shared variables and the decoder’s ability to generalize to novel conditions that it has not been trained on before.

Given that we used a total of four conditions (tactile good, tactile bad, visual good, and visual bad), there are four possible combinations available to select training and testing sets for each variable (modality and value). Similar to population decoding, we conducted CCGP on 100 iterations for each possible combination and averaged the results across iterations. Subsequently, we averaged CCGP across all four combinations and reported it as the main dataset.

### Geometric random model for testing the significance of CCGP

To assess the significance of CCGP, we followed the geometric random model used by Bernardi and colleagues^8^. This model involved creating a random geometry by relocating the cluster of points corresponding to different conditions to new random positions sampled from an isotropic Gaussian distribution. Subsequently, we rescaled all the vectors to maintain the total variance across all conditions, known as the signal variance or variance of the centroids of the clusters. These procedures allowed us to evaluate the geometry of neural patterns for each condition in a more conservative manner compared to the normal random labeling technique used in traditional population decoding.

For each possible combination of CCGP, we conducted 1000 iterations and averaged the results across combinations. Based on these averaged results for each time point, we generated a null model and tested the significance of CCGP. This rigorous approach enabled us to accurately evaluate the significance of our findings regarding the geometry of neural patterns across different conditions. The right-tailed z-test was conducted for testing statistical significance of CCGP.

### The rationale behind dividing three fixed time phases

To comprehensively examine the dynamic changes in geometries for generalized representation, we segmented neural activity into three distinct, non-overlapping phases. Phase 1 comprises pre-stimulus periods (200ms before and 150ms after the stimulus onset) to analyze neural pattern transitions before and after stimulus presentation. Phase 2, ranging from 150ms to 500ms after stimulus onset, captures the stimulus presentation period before the onset of the delay period. Phase 3 covers the entire blank delay period. These intervals were strategically chosen to optimize the visualization and analysis of neural representation geometry. We analyzed the phase 2 starting at 150 after stimulus presentation because previous research by Merchant et al. (1997)^44^ demonstrated putamen neurons begin processing tactile stimuli within the range of 100 to 200ms following stimulus onset. Furthermore, our investigation into the latency of value discrimination responses among bimodal value-coding neurons during stimulus presentation in both T-VRT and V-VRT tasks revealed median latencies of 141ms for T-VRT and 130ms for V-VRT, respectively (Hwang et el., 2024). These findings suggest that the critical period for stimulus-related information processing in the putamen occurs around the 150ms, thereby informing our phase partitioning strategy for the analysis.

### The parallelism score (PS) and its permutation test

To measure the geometry of neural representations more precisely, we estimated the Parallelism Score (PS)^8^. The weight vector derived from a trained linear classifier aligns with the vector connecting the two conditions used as the training set. In our experiment, each variable had only two coding directions from the two pairs of training conditions. We simply normalized the two weight vectors and computed the cosine of the angle between them. For example, to calculate the PS for value, we first trained two linear classifiers with [tactile good, tactile bad] and [visual good, visual bad], respectively. Then, we computed the two coding vectors from each of these classifiers and calculated the cosine angle between them. This process was repeated 100 times, and the averaged PS for each condition was reported.

To test the statistical significance of the PS, we generated 1000 number of the null model with random labeling and calculated the cosine angle of weight vectors to compare against normal PS. The two-tailed z-test was conducted for testing statistical significance of PS.

## Data and code availability

The data and codes are available from the corresponding authors on reasonable request.

## ACKNOWLEDGMENTS

This work was supported by the Korean government (MEST) grants (RS-2024-00339355) (RS-2024-00436783) through the National Research Foundation (NRF), LAMP program (RS-2023-00301976). We thank D.I. Ko for technical support and members in ILAR, SNU for technical assistance.

## Contributions

H.F.K. supervised the entire project. SH. H., JW. L., and H.F.K. performed the behavior, and single-unit recording experiments. SH. H., SP. K., JW. L., and H.F.K. analyzed the data and prepared the figures. SH. H., JW. L., and H.F.K wrote the first draft, and SH. H., JW. L., SP. K., and H.F.K. interpreted data and wrote the final manuscript.

